# A Bayesian Approach to Non-Metric Hyperbolic Multi-Dimensional Scaling

**DOI:** 10.1101/2023.12.08.570871

**Authors:** Milo Jolis, Anoop Praturu, Tatyana Sharpee

## Abstract

This paper explores the intersection of hyperbolic geometry, non-metric techniques, and Bayesian frameworks to extend the capabilities of Bayesian Hyperbolic Multi-Dimensional Scaling (HMDS). While hyperbolic geometry is gaining attention for its ability to represent hierarchical relationships, traditional metrics impose constraints on distances. Non-metric techniques offer flexibility in capturing complex structures, making them suitable for scenarios where metric distances are less meaningful. The paper introduces a novel extension of Bayesian HMDS, incorporating non-metric techniques, enabling the embedding of Euclidean data within a hyperbolic space. The approach simultaneously fits for curvature and coordinates, leveraging the scaling properties of hyperbolic space. The non-metric Bayesian Hyperbolic MDS is expected to unveil new insights into hierarchical structures within complex datasets, providing a versatile tool for analyzing high-dimensional data flexibly and accurately. The efficacy of the proposed method is demonstrated through synthetic data experiments, showcasing its ability to capture non-linear transformations and accurately predict underlying curvature, with an emphasis on its robustness to hyperparameter choices.

## 1 Introduction

In contrast to euclidean unsupervised machine learning methods, progress and development of non-euclidean techniques has been rather stagnant due to the unintuitive non-linear nature of the geometry. All be it, hyperbolic geometry is gaining support both in the machine learning and the scientific community for its predictive power and ease with which it represents complex hierarchies. Just as classical euclidean geometry can be thought of as the continuous analog of a square lattice, the same mental gymnastics can be applied to spaces of negative curvature. These spaces can be thought of the continuous analog of a tree like structure; naturally encoding hierarchy with small numbers of degrees of freedom due to the exponential expansion of the space.

Recent advances as far ranging from the studying of stress on the internet [1] to understanding place field representations in mice [2] leverage the intrinsic hierarchical nature of hyperbolic geometry. Euclidean metrics have traditionally dominated the landscape of embedding techniques and assumed linear relationships within data. While hyperbolic embeddings relieve the linearity constraint, it still imposes a strict metric relationship on distances. The limitations of metric approaches become evident when dealing with complex, non-linear structures, or when metric distances are not computable or less meaningful than a simpler dissimilarity measure. This has spurred an increased interest in non-metric techniques, which offer greater flexibility by eschewing the constraints of predefined distances. Non-metric methods excel in capturing the inherent geometry of data without assuming a linear framework, making them particularly suitable for complex structures that deviate from Euclidean norms.

The current state of non-metric techniques in the field of multidimensional scaling reflects a growing recognition of their potential [3]. While metric methods have been extensively explored and developed, the non-metric counterpart has not received commensurate attention. This underdevelopment presents an opportunity to explore new avenues and unlock the full potential of non-metric approaches.

Our focus lies at the intersection of hyperbolic geometry, non-metric techniques, and Bayesian frameworks, with a specific emphasis on extending the capabilities of Bayesian Hyperbolic Multi-Dimensional Scaling (HMDS) [4]. Hyperbolic geometry has proven advantageous in capturing hierarchical relationships, while Bayesian methodologies enhance the interpretability and uncertainty quantification of the embedding process. Building upon the foundation of metric Bayesian HMDS, our approach delves into the realm of non-metric techniques, offering a more adaptable and nuanced framework for dimensionality reduction. This novel extension not only accommodates variable curvature, allowing for the embedding of Euclidean data within a hyperbolic space, but also refines the precision of geometric parameters such as curvature and dimensionality through a Bayesian lens.

As we navigate the uncharted territory of non-metric Bayesian Hyperbolic MDS, we anticipate uncovering new insights into the hierarchical structures of complex datasets, offering a versatile tool for researchers and practitioners seeking to analyze high-dimensional data in a more flexible and accurate manner.

## 2 Non-linearity and Hyperbolic Geometry

### 2.1 Embedding Coordinates

There are many equivalent representations of hyperbolic spaces [5]. We choose to build off of the BHMDS model [4] by parameterizing our model using Lorentzian [6] coordinates. That is, on a *D* dimensional hyperbolic subspace of a *D* + 1 dimensional Minkowski space-time. Formally, our embedding coordinates are constrained to the manifold

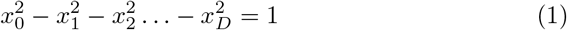

where the *D* space-like components 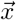 are free parameters. By solving for the time-like coordinate 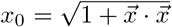 we ensure that all points lie on the future facing hyperbolic space-like manifold. Eq. (1) can be equivalently written as 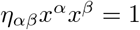 using Einstein summation where indices *α* and *β* indicate dimension and *η*_*αβ*_ = diag(1, *−*1, *−* 1, …, *−* 1) is the Minkowski metric tensor. The Lorentz model is a “native” representation of hyperbolic space where the distance to any point from the origin is just the radial coordinate *r*. The length *x* of the unique shortest path geodesic between two points who have an angular separation of Δ*θ* can be computed with the hyperbolic law of cosines.

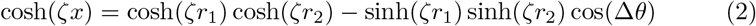

Where the curvature is defined as *K* =*− ζ*^2^. It is easy to check that in the limit of zero curvature the hyperbolic law of cosines reduces to its Euclidean form. Hyperbolic distances between any two points on the surface can now be found by combining Eq. (1) and Eq. (2) and solving for *x*.

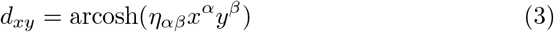

### 2.2 Scaling Curvature

It is important to note that the equations above parameterize a hyperbolic space of unit negative curvature *K* =*−*1. Here we present an algorithm that fits for curvature of the latent hyperbolic space while simultaneously addressing the non-metric nature of *δ*_*ij*_, our distance (dissimilarity) matrix. Non-metric embedding techniques are not concerned with the magnitude of input dissimilarities, caring only for their rank ordering ie. relative distances. The aim is to find a monotonic transformation *f* (*d*_*ij*_) that maps an embedding obeying metric properties *d*_*ij*_ to the non-metric space of the data. By non-linearly rescaling distances as a function of their rank, metric constraints like the triangle inequality

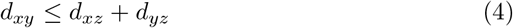

are **not** required to hold. This works quite nicely with the scaling properties of hyperbolic geometry. While traditional scale invariance observed in flat spaces does not hold, hyperbolic spaces retain a nuanced form of scale invariance.

In the context of the algorithm described in section 3, consider a rescaling of distances *d*_*xy*_→ *λd*_*xy*_. Then, according to Eq. (2), coordinates and curvature must be rescaled as *r*→ *λr* and *K*→ *λ*^*−*2^*K* respectively to preserve the original structure. From here it follows that there are two equivalent representation for a hyperbolic space up to a scaling of distance: a space with maximum radius *R*_*max*_ and unit curvature and one with *R*_*max*_ = 1 and curvature 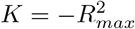.

By allowing our algorithm to fit for the coordinates 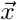 and transformation *f* (*d*_*ij*_) concurrently we exploit the scaling properties of hyperbolic space to fit for the curvature and tease out the non-metric relations in the data. Setting the embedding manifold to have unit curvature gives the points furthest from the origin freedom to expand or contract to the maximal radius of the latent space while the monotonic nonlinearity simultaneously scales these distances back.

## 3 Bayesian Non-metric HMDS

### 3.1 The likelihood function

Given a matrix *δ*_*ij*_ of distances (dissimilarities) between data points, Multi-Dimensional Scaling seeks an embedding of points 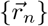 in a geometric space whose distance matrix *d*_*ij*_ matches the dissimilarity matrix as closely as possible [7]. This is formulated by defining a stress function which is minimized when the matrices are exactly equal:

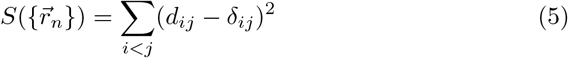

One can obtain the embedding 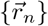 through straightforward gradient descent on *S*. Non-metric MDS extends this formulation by additionally fitting for a monotonic transformation *f* which allows for the matrix of data dissimilarities *δ*_*ij*_ to violate the metric conditions.

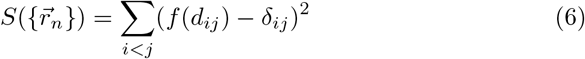

As in [4] we formulate MDS with a generative stochastic model [8] instead of in terms of a stress function. We assume the data dissimilarities are generated directly from an underlying geometric space by a process which monotonically transforms then and adds white noise to the system

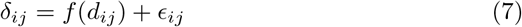

where *ϵ*_*ij*_ *∼ 𝒩* (0, *σ* _*i j*_) are independent normally distributed noise and *f* is an arbitrary monotonic transformation. As in [4] we assign an uncertainty *σ*_*i*_ to each embedded point and infer the uncertainty of the distance between *i* and *j* as 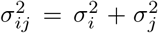. From this we can easily write the likelihood of a single observation as

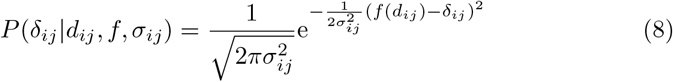

And thus the negative log-likelihood of the entire data matrix is given by

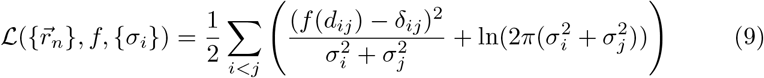

### 3.2 The posterior distribution

To complete our model we need to specify how we intend to infer the proper monotonic function, and place priors on our parameters in order to regularize them.

Since the space of all possible *f* is infinite dimensional, we must find a low dimensional parameterization of *f* in order to fit for it. We do so with a series of step-like functions defined as follows. We define a unit step *ϕ*_*b*_(*x − s*) as

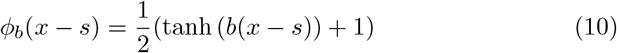

This is a unit sized step function located at *x* = *s* whose slope at *s* is *b/*2. We define *f* as a sum of *N*_*s*_ steps of varying heights

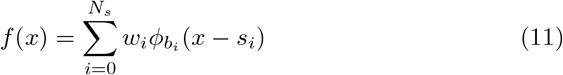

The 3*N*_*s*_ parameters {*w*_*i*_, *b*_*i*_, *s*_*i*_} this completely determine the monotonic transformation and can be fit for. Note that all of these parameters are positive. In practice we find that we only need a few step functions to accurately fit a wide variety of transformations, which we study in more detail in later sections.

Placing inverse normal priors in the uncertainty parameters and half normal priors on the step function parameters we can write down the full loss function as the negative log posterior

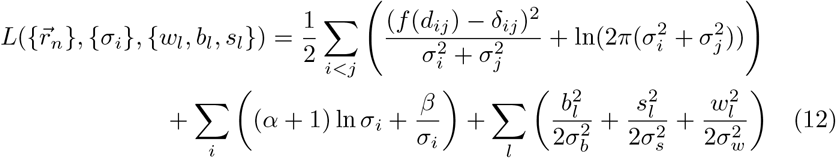

Similar to the approach in [4] we employ the L-BFGS algorithm from [9], a statistical programming language distributed under a BSD license, to minimize Equation (12).

## 4 Synthetic Data Results

### 4.1 Fitting the coordinates and transform

Initially, we assess our approach using synthetic distance matrices and complex monotonic functions to verify that the algorithm fits the nonlinearity well. Mock distance matrices are generated by uniformly sampling 100 points in hyperbolic spaces of variable dimension up to a specified maximum radius. These matrices are then normalized such that max(*δ*_*ij*_) = 1. This ensures that our algorithm has no knowledge about the curvature of the hyperbolic space that generated the data. Next, iid. noise of magnitude 0.05*R*_*max*_ is added to mimic real datasets. The complex nonlinearity *t*(*x*) is randomly generated by sampling 25 frequencies and amplitudes from normal distributions *f ∼ 𝒩*(0, 10) and *A ∼ 𝒩* (0, 1) respectively. These values are passed into *A*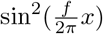 functions which are summed and then cumulatively integrated. We normalized the integral such that *t*(*x*) : [0, 1] →[0, 1]. Each distance is passed through the complex monotonic function to create our non-metric dissimilarity matrix then see how well our algorithm can reproduce the transformation. We find that setting the initial conditions of *f* to be the identity transform helps the algorithm converge to its global optima. Fig. 1 shows the quality of fits for 50 points sampled from *R*_*max*_ = 5 and *D* = 3. The embeddings were done in the same dimension.

**Figure 1:**
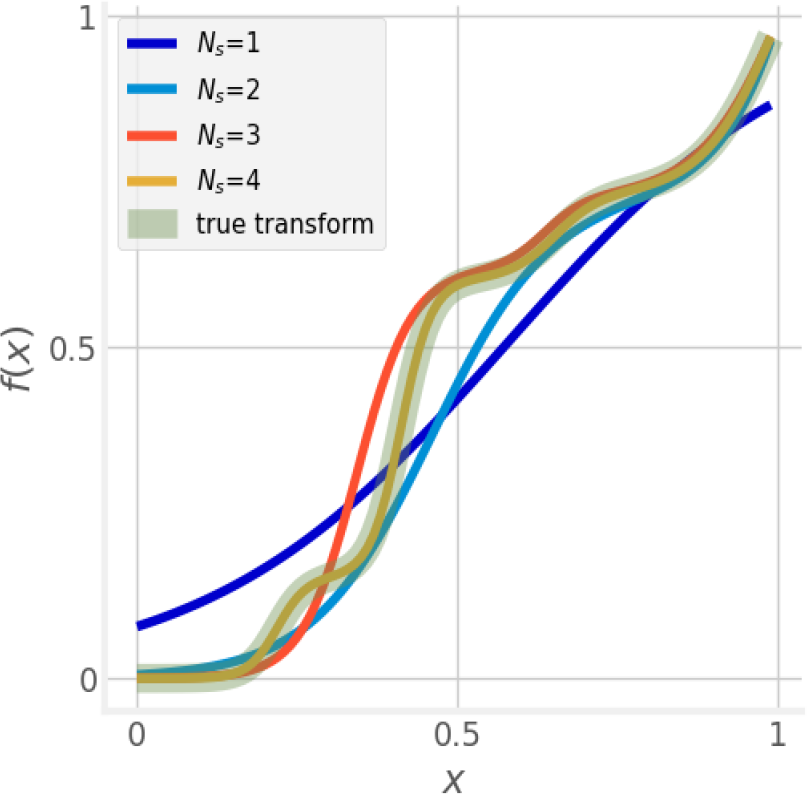
Comparing the fits using different values of *N*_*s*_ to the true transform.

**Figure 2:**
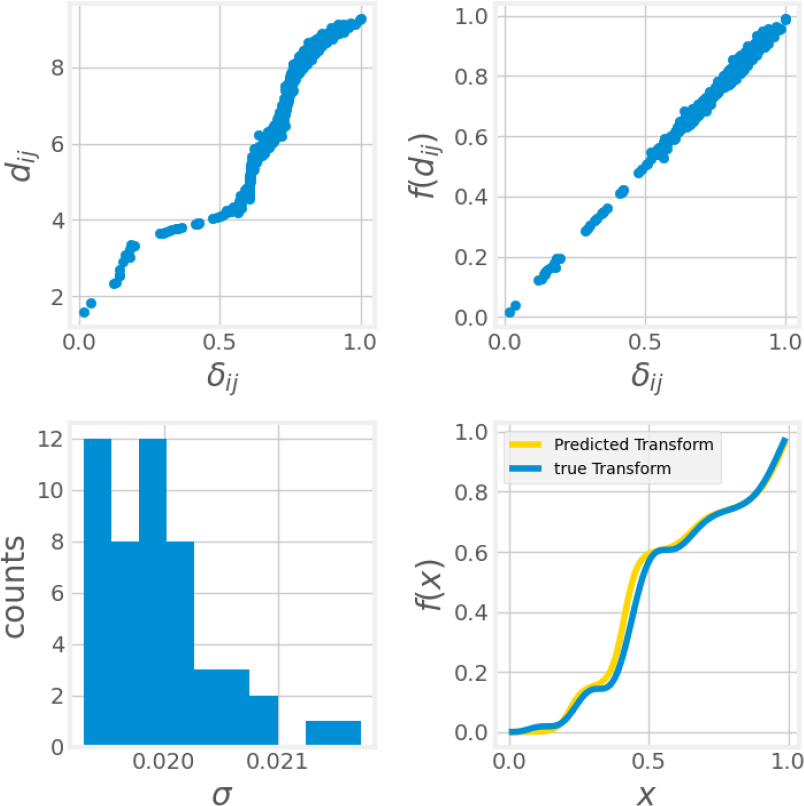
Top Left: Shepard diagram of metric embedding distances versus non-metric data distances. Top Right: Shepard diagram after fitted transform is applied to embedding distances. Bottom Left: Distribution of embedding uncertainty parameters. Bottom Right: Comparison of true transformation and fitted transformation.

It is quite apparent that the best fit produced by the algorithm is for *N*_*s*_ = 4, however, the other fits for different numbers of *N*_*s*_ are still quite good. We elaborate on this in Sec. 4.2.

As shown, our non-metric MDS algorithm is capable of reproducing the monotonic transformation of distances quite well. This, however does not mean that the actual fit of the data points was performed correctly. To illustrate that the algorithm approximately reproduces the input distances for the *N*_*s*_ = 4 embedding, we show the shepard diagram of the input distances plotted against the transformed embedding distances. For the same embedding we also show a histogram of the pointwise embedding uncertainties *σ*. Both of these are indicators to whether the embedding was successful.

The Shepard diagram shows a very high correlation of the input disimilarities with the transformed embedding distances and the histogram of uncertainty values shows that the uncertainty of the position of the points was very low, both indicators that embedding was fit correctly.

### 4.2 Fitting for dimension and number of steps

We’ve shown that our non-metric hyperbolic MDS algorithm can correctly embed points that are related through non-metric dissimilairities by finding a monotonic nonlinear scaling of the embedding distances concurrently. Now we turn our attention to the hyperparameters of the method. One of the best use cases for MDS is finding a low dimensional representation of the original dataset. However one must be certain that the embedding is the optimal one in order to extract meaningful information from it. Since we are working with a Bayesian framework we can actually formulate a way to find the optimal hyperparameter of the embedding. This is done by computing the evidence [10]. The Bayesian Information Criterion (*BIC*), integrates out the size effects of the likelihood ***ℒ***_***D***_**(*θ*)**of a *D* dimensional embedding or 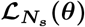of an embedding with *N*_*s*_ steps. Here ***θ***denotes the model parameters.

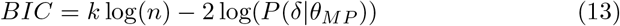

Here, *k* denoted the number of parameters and *n* the number of observations*θ*_*MP*_ are the parameters of the final embedding assuming a global optima was reached and the posterior was maximized.

A random distance matrix of 50 points was generated using the same method as in Sec. 4.1 with *R*_*max*_ = 2.5 and *D* = 5. It was then processed with 10 different random monotonic transformations and each resulting matrix was fit to 9 different dimensions 8 times, varying the number of steps in the fit for monotonic transformation. The *BIC* was computed for every embedding and the resulting averages and standard deviation of dimension *D* and number of steps *N*_*s*_ are ploted in Fig. 3.

**Figure 3:**
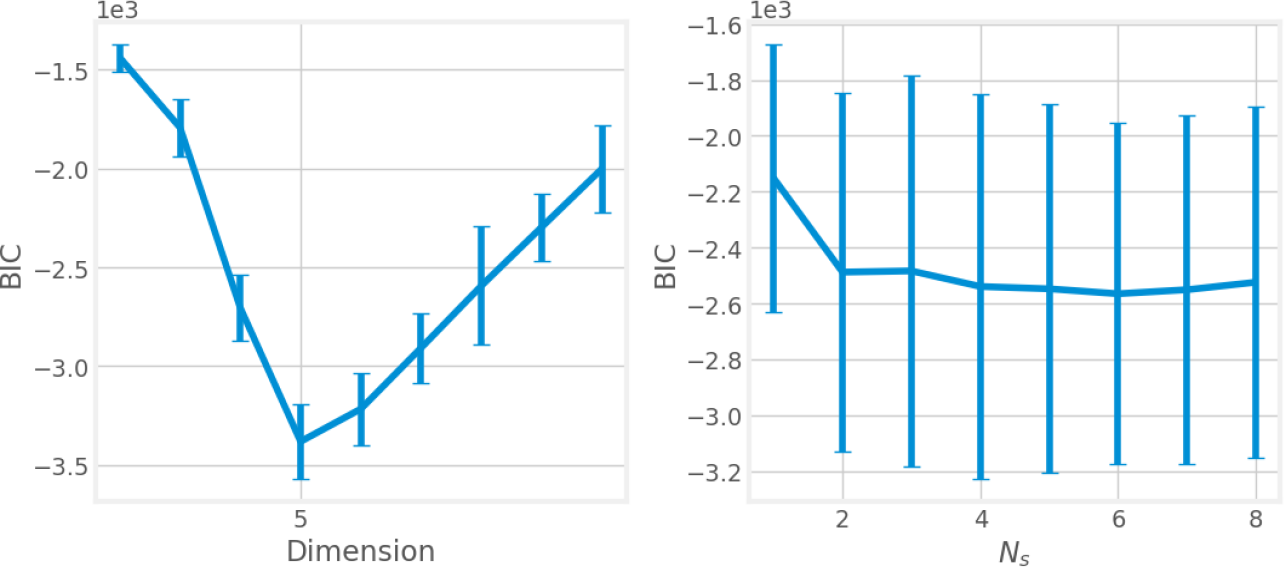
Left: BIC as a function of dimension correctly predicts underlying dimension of data. Right: BIC shows that model is insensitive to number of step parameters used.

The model has very high predictive power over the correct embedding dimension as shown in the figure. There is a clear minimum in the *BIC* at *D* = 5. As we increase the dimension of the embedding space starting at *D* = 2 the model is able to better fit the for the true non-metric distances. When it passes *D* = 5 then the fit is just as good but the model is penalized for the unecesairy degrees of freedom. The model shows high evidence for the true dimensionality of the data but less so over the number of steps. This is likely to have been caused by the fact that all of the generated monotonic transforms were random and hence had random number of steps within them causing an averaging effect in the *BIC*. However, under the assumption that the non-metric data has a true latent monotonic nonlinear rescaling of distances then we expect this Bayesian framwork to uncover it. It must also be noted that in Fig. 1, all of the fitted monotonic functions that were not optimal still provided a good estimate to the true one. We interpret this finding as the model performing well even when the number of steps in the monotonic fitted functions is not the base truth.

### 4.3 Fitting for curvature

Finally, to fit for curvature we generated 3 different distances matrices of 100 points each from hyperbolic spaces of *D* = 3 and varying curvature *R*_*max*_ =[0.1, 0.5, 1, 2, 3, 4, 5, 6, 7, 8, 9, 10]. Every distance matrix was subject to 2 different transformations and they were each fit for *N*_*s*_ = [1, 2, 3, 4, 5].

We see from figure 4 that the model is able to successfully predict the curvature for a wide range of simulated data. We do however see that the model is not optimal at extremely low curvatures and extremely high curvatures. For the low curvature limit we see in the second panel of figure 4 that the uncertainty parameters of those embeddings are quite high, so we attribute the model’s failure to match the curvature to poor fitting. In the high curvature limit, the problems arise from fundamental difficulties with hyperbolic geometry. Since the volume of the space grows exponentially with curvature and radius the majority of the space is dominated by points near the outer radius. Thus we see significant compression in the distribution of distances concentrating around *d* = 2*R*. Further compression of the distance matrix from the non-linear transformation makes curvature inference extremely difficult in this limit since only a small window of possible distances are being sampled from. We plan on investigating non-uniform distributions of points in the high curvature limit to see if we can remedy this fundamental difficulty.

**Figure 4:**
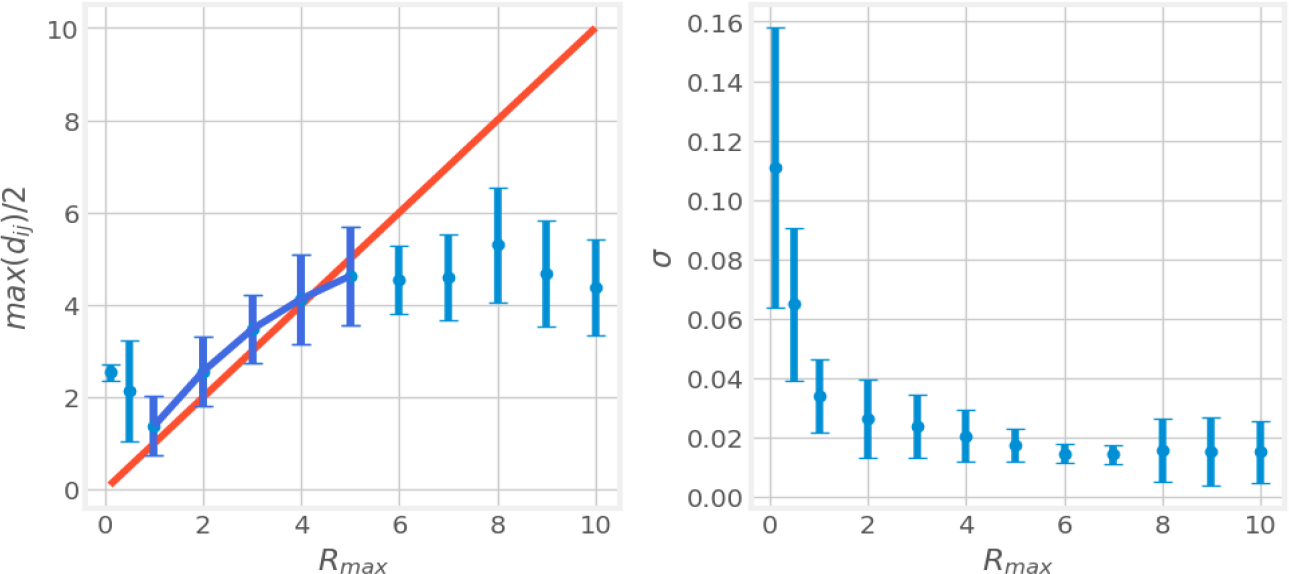
Left: Sampling radius (predicted curvature) as a function of true curvature. Right: Uncertainty parameters of each embedding.

